# Exploring the impact of digestive physicochemical parameters of adults and infants on the pathophysiology of *Cryptosporidium parvum* using the dynamic TIM-1 gastrointestinal model

**DOI:** 10.1101/2024.07.17.603888

**Authors:** Julie Tottey, Lucie Etienne-Mesmin, Sandrine Chalançon, Alix Sausset, Sylvain Denis, Carine Mazal, Christelle Blavignac, Guillaume Sallé, Fabrice Laurent, Stéphanie Blanquet-Diot, Sonia Lacroix-Lamandé

## Abstract

**Background:** Human cryptosporidiosis is distributed worldwide, and it is recognised as a leading cause of acute diarrhoea and death in infants in low- and middle-income countries. Besides immune status, the higher incidence and severity of this gastrointestinal disease in young children could also be attributed to the digestive environment. For instance, human gastrointestinal physiology undergoes significant changes with age, however the role this variability plays in *Cryptosporidium parvum* pathogenesis is not known. In this study, we analysed for the first time the impact of digestive physicochemical parameters on *C. parvum* infection in a human and age-dependent context using a dynamic *in vitro* gastrointestinal model.

**Results:** Our results showed that the parasite excystation, releasing sporozoites from oocysts, occurs in the duodenum compartment after one hour of digestion in both child (from 6 months to 2 years) and adult experimental conditions. In the child small intestine, slightly less sporozoites were released from excystation compared to adult, however they exhibited a higher luciferase activity, suggesting a better physiological state. Sporozoites collected from the child jejunum compartment also showed a higher ability to invade human intestinal epithelial cells compared to the adult condition. Global analysis of the parasite transcriptome through RNA-sequencing demonstrated a more pronounced modulation in ileal effluents compared to gastric ones, albeit showing less susceptibility to age-related digestive condition. Further analysis of gene expression and enriched pathways showed that oocysts are highly active in protein synthesis in the stomach compartment, whereas sporozoites released in the ileum showed downregulation of glycolysis as well as strong modulation of genes potentially related to gliding motility and secreted effectors.

**Conclusions:** Digestion in a sophisticated *in vitro* gastrointestinal model revealed that invasive sporozoite stages are released in the small intestine, and are highly abundant and active in the ileum compartment, supporting reported *C. parvum* tissue tropism. Our comparative analysis suggests that physicochemical parameters encountered in the child digestive environment can influence the amount, physiological state and possibly invasiveness of sporozoites released in the small intestine, thus potentially contributing to the higher susceptibility of young individuals to cryptosporidiosis.

## BACKGROUND

The zoonotic disease cryptosporidiosis caused by the protozoan Apicomplexa parasite *Cryptosporidium* is endemic in many low-income countries and potentially epidemic in high-income countries. This enteric disease leads to watery diarrhoea that can be life-threatening in individuals with an immature or a compromised immune system. Results from cohort studies have consistently shown that young age is associated with a higher risk of *Cryptosporidium* infection (1). Accordingly, *Cryptosporidium* infection is particularly associated with prolonged (7–14 days) or even persistent (≥ 14 days) diarrhoea during childhood, and can also lead to malnutrition and growth deficiency (1). Cryptosporidiosis is considered as one of the most common causes of infectious moderate-to-severe diarrhoea in children under the age of two (2,3) and an important contributor to early childhood mortality in low-resource settings (4), thus constituting a serious public health concern. In 2016, acute infections due to *Cryptosporidium* caused more than 48000 deaths in children under 5 years and more than 4.2 million disability-adjusted life-years lost (5).

*Cryptosporidium parvum* is one of the two most relevant *Cryptosporidium* species to humans and is transmitted primarily by the fecal–oral route either by direct contact with an infected human or animal or indirectly via food or water contaminated by oocysts (6). Once ingested, *C. parvum* oocysts excyst in the gastrointestinal tract, releasing infective and motile sporozoites that invade intestinal epithelial cells. The parasite life cycle then progresses through three rounds of asexual replication (7) before shifting to sexual reproduction and then to the production of a high number of infectious oocysts that ensure either auto-infection of the same host by infecting nearby intestinal cells or transmission in the environment after release in the faeces.

The higher incidence and severity of cryptosporidiosis reported in children under the age of two can be first attributed to their immature immune status, making them as susceptible to the infection by the parasite as immunocompromised adults (8). Nevertheless, other factors, such as those associated with the digestive environment (*i.e.,* immaturity of digestive processes, intestinal epithelium and resident microbiota) where *C. parvum* infection takes place, could also contribute to the age-dependent nature of symptomatic cryptosporidiosis. Consequently, there is a crucial need to investigate *C. parvum* pathogenesis considering the gastrointestinal physiology that varies greatly with age (9), especially in light of the differences in digestive physicochemical parameters encountered in children, compared to adults.

Due to obvious ethical and technical reasons, it remains very difficult to evaluate the pathophysiology of food- or water-borne pathogens in the human gastrointestinal tract, especially in pediatric populations. *In vitro* gastrointestinal models represent an appropriate alternative to *in vivo* assays to study the impact of digestive physicochemical parameters alone (*i.e.,* independently of any other influencing factors from the host) in a nutritional, pharmaceutical, toxicological or microbiological context. Among the available gastrointestinal systems, the computer-controlled TNO (Toegepast Natuurwetenschappelijk Onderzoek) gastroIntestinal Model-1 (TIM-1), which combines multi-compartmentalisation and dynamism, is one of the most complete *in vitro* simulators of the human upper gastrointestinal tract (10). By reproducing physiologically relevant conditions, the TIM-1 system allows the closest simulation of *in vivo* dynamic events occurring within the human stomach and three compartments of the small intestine. As a result, it has been successfully used for a diversity of applications in the past, for instance in an array of pathogen-related microbiological studies (10–16). As an example, the TIM-1 model has been used to demonstrate that the variability in human digestive physicochemical parameters that occurs between child and adult populations could play a role in *E. coli* O157:H7 pathogenesis, which is considerably more severe in children than in adults (12).

Within this framework, we designed an original approach to question for the first time the impact of digestive physicochemical parameters on *C. parvum* infection, in a human and age-dependent context. The aim of the present study was to use the sophisticated TIM-1 model for a comparative analysis of *C. parvum* infection under the digestive conditions encountered in young children (from 6 months to 2 years) or in adults following the simulated digestion of a glass of contaminated water. Various parasite parameters, from the parasite excystation kinetics and global gene expression, to the sporozoite physiological state and infectivity, were monitored throughout the simulated human digestions.

## METHODS

### Cells and parasites

Human ileocecal adenocarcinoma cells (HCT-8) were purchased from the American Type Culture Collection, cultured in RPMI 1640 with phenol red supplemented with 2 mM GlutaMAX™, 10% (v/v) heat-inactivated fetal bovine serum (FBS), 1 mM sodium pyruvate, 50 U/ml penicillin, and 50 µg/ml streptomycin and maintained at 37°C in a humidified atmosphere under 5% CO_2_. The *C. parvum* INRAE Nluc strain was generated by transgenesis to produce Nluc-expressing parasites, and purified as described in Swale *et al.*, 2019 (17).

### *In vitro* digestions in the TIM-1 gastrointestinal model

The TIM-1 model (TNO, Zeist, The Netherlands) consists of four successive compartments simulating the human stomach and the three segments of the small intestine (duodenum, jejunum, and ileum) (Figure 1). The main parameters of digestion, such as body temperature, pH, peristaltic mixing and transport, gastric, biliary, and pancreatic secretions and passive absorption of small molecules and water, are reproduced as accurately as possible, as already described (16). The TIM-1 system was programmed to reproduce, based on *in vivo* data, the physicochemical digestive conditions observed in a healthy adult or a young child (from 6 months to 2 years) when a glass of water is ingested as previously described (12). The TIM-1 system was inoculated with a parasite suspension consisting of 200 mL of mineral water (Volvic®, Danone, France) experimentally contaminated with 1 × 10^8^ oocysts of the *C. parvum* INRAE Nluc strain. Two sets of experiments were performed: gastric digestions where the stomach compartment was solely used (total duration of 60 min) and gastrointestinal digestions using the entire TIM-1 model (total duration of 300 min). Digestions were run in triplicate.

**FIGURE 1.**
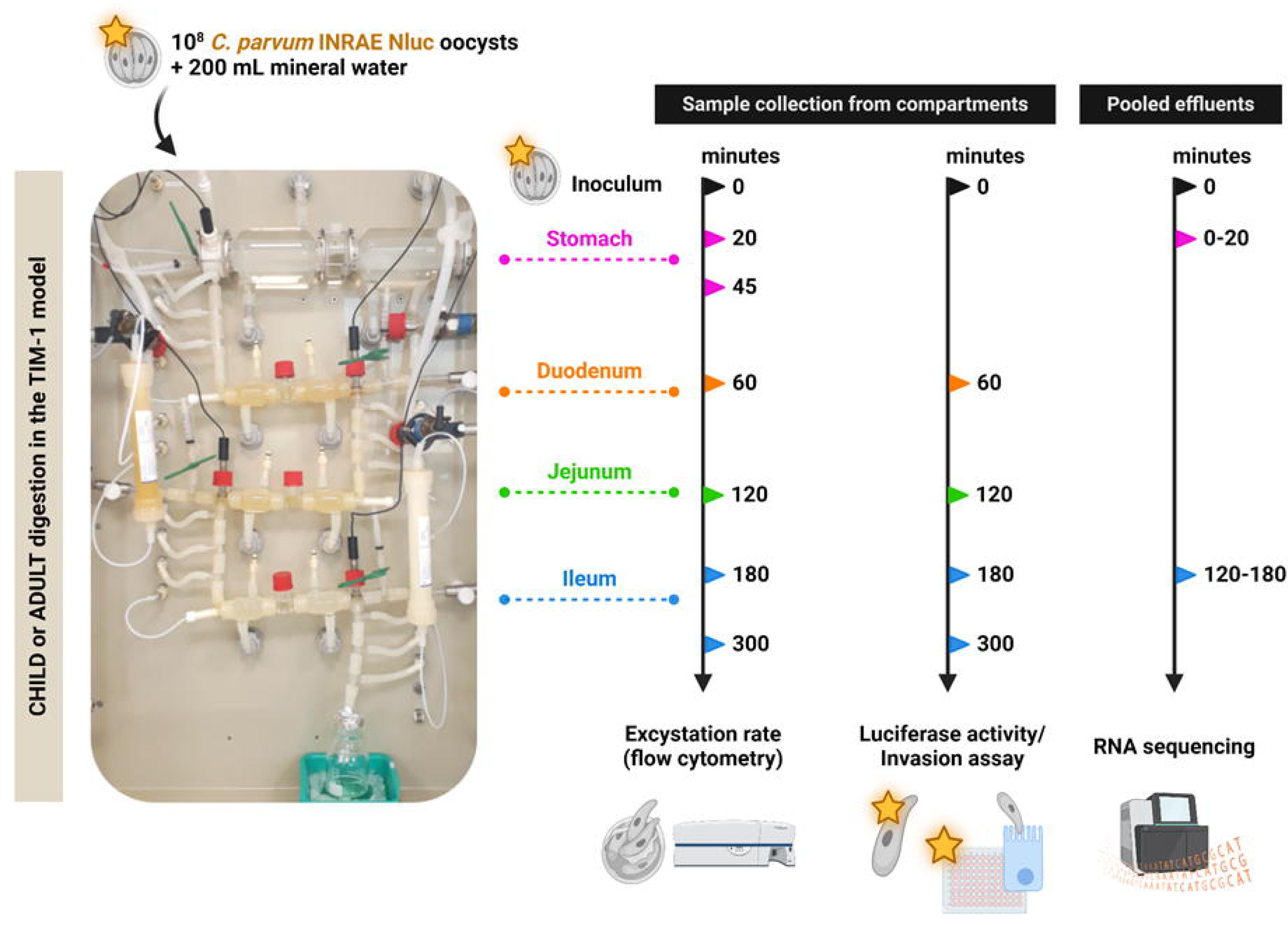
TIM-1 experimental set-up and analysis. The left side shows a picture of the TIM-1 system composed of four successive compartments simulating the human stomach and the three parts of the small intestine (*i.e.,* the duodenum, jejunum and ileum) mimicking the main physicochemical parameters of human gastrointestinal digestion. Digestion experiments were performed to reproduce an adult or an infant (from 6 months old to 2 years old) ingesting a glass of water contaminated with 1 × 10^8^ oocysts of the *C. parvum* INRAE Nluc strain. On the right side, sampling times (minutes) are indicated when samples were taken either directly from each compartment (for flow cytometry, sporozoite luciferase activity and invasion assay); or indirectly by pooling the gastric effluents when the stomach compartment was solely used, or the ileal effluents when the entire TIM-1 system was used (for RNA sequencing). Digestions were run in triplicate. Created with BioRender.com (Agreement number: JH26ZFM4TU).

### TIM-1 sampling

Samples were taken in the initial parasite suspension (t=0) and regularly collected during *in vitro* digestions from each digestive compartment (stomach, duodenum, jejunum, and ileum) to evaluate parasite excystation kinetics, sporozoite luciferase activity and host cell invasion (Figure 1). Gastric and ileal effluents were also collected on ice and pooled on 0-20, 20-45, and 45-60 min for gastric digestions and hour-by-hour during 5 h for gastrointestinal digestions. Gastric and ileal effluents collected during 0-20 min and 120-180 min, respectively, were used for RNA extraction.

### Parasite excystation kinetics

The parasite excystation success was monitored in three independent experiments by flow cytometry from samples collected regularly from each digestive compartment during *in vitro* digestions in the TIM-1 system. Prior to each TIM-1 digestion assay, flow cytometry controls and gating strategy on forward-angle light scatter/side-angle light scatter were performed based on analysis of intact (non-excysted) oocysts, and non-filtered or 5 µm-filtered (Sartorius AG, Göttingen, Germany) *in vitro* excysted parasites (Figure 2.A). Such controls aimed to differentiate *C. parvum* oocysts from sporozoites and from the background. Flow cytometry analyses were all performed immediately after sample collection on a BD™ LSR II cytometer and data were collected with the BD FACSDiva™ Software version 9 (BD Biosciences, Franklin Lakes, USA). Results are expressed as relative percentages of parasites detected as intact oocyst stages in each digestive compartment compared to the corresponding age condition’s inoculum.

**FIGURE 2.**
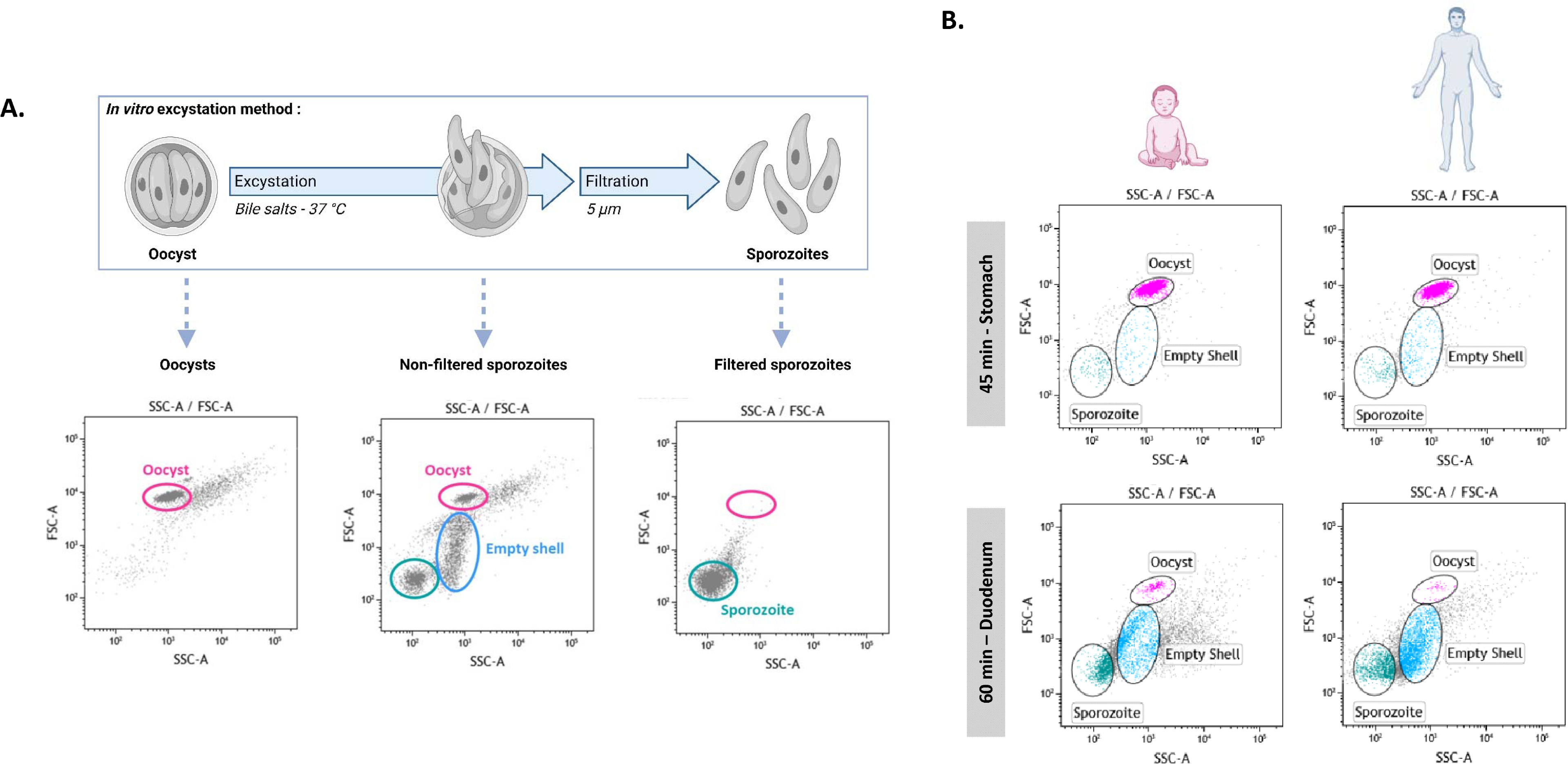
*Cryptosporidium parvum* excystation in the human *in vitro* upper gastrointestinal tract. (A) Flow cytometry gating strategy on forward-angle light scatter/side-angle light scatter established during *in vitro* excystation of parasites, allowing detection of intact oocysts, empty oocyst shells and sporozoites. Created with BioRender.com (Agreement number: FF26ZFMVSM). **(B)** Parasite excystation during child or adult digestion in the TIM-1 system. Cytograms obtained for one representative experiment for the child or the adult digestion at 45 min in the stomach and at 60 min in the duodenum show the timing when excystation occurs in the *in vitro* gastrointestinal tract.

### Sporozoite luciferase activity

The physiological state of *C. parvum* sporozoites was monitored in three independent experiments for the initial parasite suspension and for samples collected regularly from the duodenum, jejunum and ileum compartments during *in vitro* digestions in the TIM-1 system. Immediately after collection, each sample was washed with phosphate-buffered saline (PBS), filtered through a 5 µm membrane (Sartorius AG, Göttingen, Germany) to discard oocysts and empty shells, centrifuged (10000 ×g, 3 min, 4°C) and resuspended in PBS. The luciferase activity expressed by the recovered sporozoites was assessed (three replicates for each sample) using the Nano-Glo® Luciferase Assay System (Promega Corporation, Madison, USA). The sporozoites were mixed (v/v) with the Nano-Glo® Luciferase Assay Buffer containing 1:50 of the Nano-Glo® Luciferase Assay Substrate and the luminescence was measured with the GloMax®-Multi Detection System (Promega Corporation, Madison, USA). The luciferase activity expressed by sporozoites collected from the TIM-1 compartments was normalised to the initial parasite suspension for each assay.

### Parasite invasion assay

The ability of *C. parvum* sporozoites to invade intestinal epithelial cells was monitored in two independent experiments for samples collected regularly from the small intestinal compartments during *in vitro* digestions. Immediately after collection, each parasite sample was washed with sterile PBS, filtered through a 5 µm membrane (Sartorius AG, Göttingen, Germany) to discard oocysts and empty shells, centrifuged (10000 ×g, 3 min, 4°C) and resuspended into sterile PBS. The recovered sporozoites were immediately used to infect HCT-8 cell monolayers grown to 80% confluence in Nunc™ F96 MicroWell™ white polystyrene plates (Thermo Fisher Scientific, Waltham, USA). After 2.5 hours, cells were washed twice with sterile PBS and used to analyse the invasion of parasites (six replicates per sample). The luciferase activity was detected directly from the infected cell monolayers using the Nano-Glo® Luciferase Assay System (Promega Corporation, Madison, USA) and the GloMax®-Multi Detection System as described above. Luciferase activity data were normalised to the luminescence background detected in uninfected cultures.

### Parasite RNA extraction

RNA samples were collected on ice from the initial parasite suspension and from gastric (0-20 min) or ileal (120-180 min) effluents during *in vitro* digestions. Samples were then centrifuged (10000 ×g, 5 min, 4°C), resuspended into 500 μL TRI Reagent® (Sigma-Aldrich, Saint-Louis, USA) and subjected to 5 cycles of [1 min vortexing - 1 min on ice] after addition of 0.5 mm glass beads. The lysates were centrifuged (10000 ×g, 10 min, 4°C) and the supernatants were mixed with absolute ethanol and transferred to a Zymo-Spin™ IC column (Direct-zol™ RNA Microprep kit, Zymo Research, Irvine, USA). Total RNAs were isolated according to the manufacturer’s instructions. Any contaminating genomic DNA was removed using the DNase I Set kit (Zymo Research). RNA was further purified with the RNA Clean & Concentrator™-5 kit (Zymo Research) and concentrations measured using the Qubit™ 2.0 fluorometer using RNA HS assay kit (Thermo Fisher Scientific). RNA quality was assessed using the Agilent 2100 Bioanalyzer system and the RNA 6000 Pico kit (Agilent Technologies, Santa Clara, USA).

### RNA-Seq analysis of differentially expressed genes

Library construction and sequencing were performed by Helixio (Biopôle Clermont-Limagne, Saint-Beauzire, France). The RNA library preparation was performed using the QuantSeq 3′ mRNA-Seq Library Prep Kit FWD for Illumina (Lexogen, Vienna, Austria). A total of 15 RNA-Seq libraries were generated, corresponding to 3 libraries for each experimental group, and then sequenced with the NextSeq® 500 System (Illumina, San Diego, USA) using a single-read sequencing of 76 bp length configuration. Raw sequence data were assessed for quality using the FastQC v0.11.3 tool (Babraham Institute, Cambridge, UK). Sequenced reads were aligned to the *C. parvum* IOWA-ATCC reference genome (genome assembly ID: ASM1524537v1) using the STAR software (18). The number of sequences reads that mapped the *C. parvum* genome varied between 11 and 14 million per library. Gene counts were determined using the STAR software. The R–based packages DESeq2 (19) and edgeR (20) were used to normalise data, perform descriptive analysis as well as all pairwise comparisons to determine the differentially expressed genes between experimental groups. Clustering of transcriptomic profiles was assessed through a principal-component analysis (PCA) using normalised RNA-Seq data of a set of 2430 filtered genes, for which at least 10 reads were counted in a minimum of three samples. The *P*-values generated during pairwise comparisons were adjusted for multiple testing with the Benjamini-Hochberg procedure which controls for false discovery rate (FDR). Genes were considered to be differentially expressed between two experimental groups at an FDR adjusted *P*-value < 0.05. Gene Ontology (GO) (21,22) and KEGG Metabolic Pathway (23–25) enrichment analyses were performed on differentially expressed genes on CryptoDB (26).

### Statistical analysis

Graphs and boxplots were generated using GraphPad Prism version 10.2.3 and statistical analyses were performed using R version 4.3.1. Significant differences in luciferase activity and invasion ability data according to time of digestion were tested using a nonparametric analysis of longitudinal data with the R package “nparLD” (27) version 2.2. In case of a significant time effect, pairwise comparisons between time points were calculated with the R package “nparcomp” (28) version 3.0. *P* < 0.05 was considered statistically significant and indicated by non-corresponding letters (lowercase for the child condition and capitals for the adult condition). The Mann-Whitney non-parametric test was performed to test the effect of treatment (child *vs.* adult) for each time point and to analyse luciferase data collected from live or heat-killed parasites (Additional file 1), with significant differences indicated as follows: * *P* < 0.05; ** *P* < 0.01; *** *P* < 0.001; **** *P* < 0.0001.

## RESULTS

### Parasite excystation kinetics in the simulated upper gastrointestinal tract

To establish the gating strategy, flow cytometry controls were performed during *in vitro* excystation of parasites (Figure 2.A). These controls showed that intact oocysts were abundant before excystation and easily discriminated from empty shells and sporozoites during the process of excystation (when non-filtered). In contrast, intact oocysts were scarcely detected in filtered excysted samples, in which mostly sporozoites could be observed.

The same gating strategy was then used to monitor parasite excystation kinetics during child and adult digestions in the TIM-1 system. Time of digestion had an overall significant effect on parasite excystation in both child (*P* < 0.0001) and adult (*P* < 0.0001) conditions. In the simulated stomach compartment, the proportion of intact oocysts (*i.e.,* non-excysted parasites) remained high regardless of the age condition, with 80.4 ± 7.2% and 81.4 ± 10.1% intact oocysts detected at 45 min for child and adult digestion, respectively (Figure 2.B, Table 1). In contrast, subsequent transit in the duodenal compartment marked a brutal decrease in the proportion of intact oocysts detected, which dropped to 4.5 ± 0.3% and 1.5 ± 0.3% at 60 min for child and adult condition, respectively. The parasite excystation further progressed in the small intestine compartments and reached a minimal amount of intact oocysts detected for both conditions in the ileal compartment by the end of digestion (1.0 ± 0.1% and 0.6 ± 0.1% at 300 min for the child and the adult digestion, respectively). The quantity of intact oocysts detected in the child’s small intestine was consistently higher when compared to the adult condition whatever the compartment and time considered (Table 1). Our results suggest that sporozoites released from excystation in the small intestine would be slightly less abundant in infants compared to adults.

**Table 1.** Percentage of parasites detected as intact oocysts for the child or adult TIM-1 experiments. The parasite excystation kinetics was analysed by flow cytometry during child or adult digestion in the TIM-1 system. Samples recovered from the inoculum (Ino) or from the stomach (Sto), duodenum (Duo), jejunum (Jej) or ileum (Ile) compartments were immediately processed after collection. Values from three independent child or adult digestion in the TIM-1 system are expressed as relative percentages of intact oocysts as compared with that of the inoculum for each digestion assay.

| Age condition | TIM-1 digestion | 0 min Ino | 20 min Sto | 45 min Stomach | 60 min Duo | 120 min Jej | 180 min Ile | 300 min Ile |
| --- | --- | --- | --- | --- | --- | --- | --- | --- |
| CHILD | 1 | 100 | 98.2 | 93.5 | 3.8 | 1.9 | 1.8 | 1.1 |
|  | 2 | 100 | 73.5 | 68.6 | 4.7 | 2.6 | 2.4 | 0.9 |
|  | 3 | 100 | 86.8 | 79.0 | 5.0 | 2.7 | 2.6 | 1.0 |
| ADULT | 1 | 100 | 101.5 | 99.5 | 1.4 | 1.3 | 1.2 | 0.9 |
|  | 2 | 100 | 83.2 | 64.7 | 2.0 | 1.4 | 1.1 | 0.6 |
|  | 3 | 100 | 90.1 | 79.9 | 1.0 | 0.7 | 0.6 | 0.4 |

### Physiological state and invasion ability of *C. parvum* sporozoites collected from the *in vitro* small intestine

The physiological state of the sporozoites released following excystation in the small intestine of the TIM-1 system was estimated through analysis of the activity of the luciferase reporter gene that is constitutively expressed by the parasite strain (Figure 3.A). Time of digestion had an overall significant effect on sporozoite luciferase activity in both child (*P* < 0.0001) and adult (*P* < 0.0001) conditions. In the child small intestine, sporozoite relative luciferase activity significantly increased from 38.9 ± 12.1 normalised relative light units (RLUs) in the duodenum at 60 min and 950.5 ± 12.1 RLUs in the jejunum at 120 min to a maximum reached at 180 min in the ileum (2.0 × 10^4^ ± 4.2 × 10^3^ RLUs; *P* < 0.0001 between 60 min and 180 min and between 120 min and 180 min). During the last two hours of digestion, the relative luciferase activity detected in the child ileal compartment decreased (7.9 × 10^3^ ± 1.4 × 10^3^ RLUs) but remained significantly higher than the one detected in the duodenum and jejunum at the beginning of digestion (*P* < 0.0001 between 60 min and 300 min and between 120 min and 300 min). The luciferase activity kinetics of sporozoites collected from the adult digestions followed the same trend but at a much lower intensity, with a maximum reached of 706.7 ± 80.0 RLUs at 180 min in the ileum. Additionally, the relative luciferase activity expressed by sporozoites was consistently and significantly higher for the child condition when compared to the adult one, whatever the compartments of the small intestine and the time points (*P* < 0.0001 at 60, 120, 180 and 300 min). Controls performed on the *C. parvum* INRAE Nluc strain showed that the luciferase activity measured from excysted parasites was dependent on the quantity (*P* < 0.01 between 1 × 10^7^ and 1 × 10^8^ excysted parasites) and viability (*P* < 0.01 between live and heat-killed parasites) of parasites present in the sample (Additional file 1). In the TIM-1 small intestine under child condition, fewer sporozoites (Figure 2.B, Table 1) but a significantly higher luciferase activity (Figure 3.A) was detected compared to the adult condition.

**FIGURE 3.**
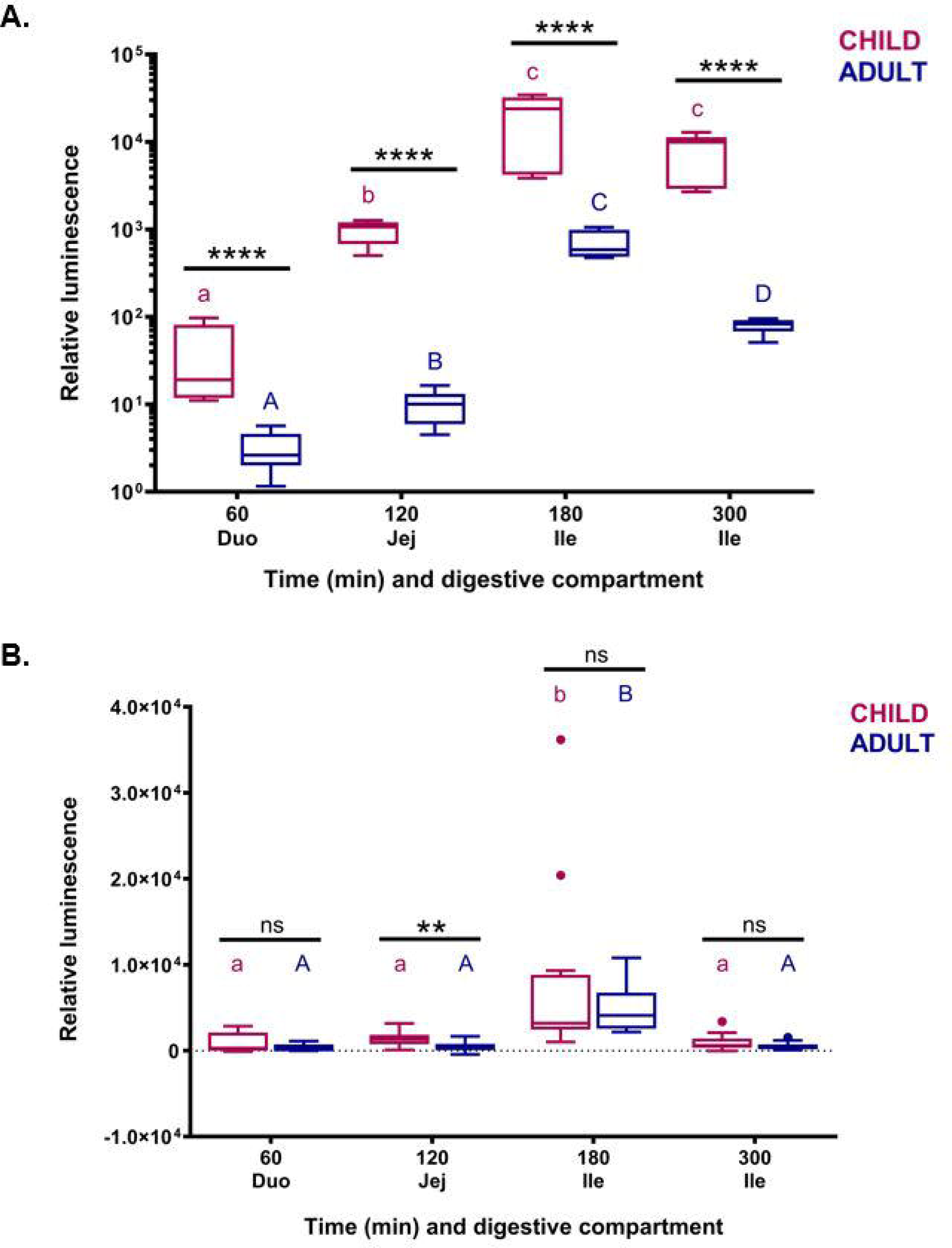
Physiological state and invasion ability of *Cryptosporidium parvum* sporozoites collected from the TIM-1 system. **(A)** Luciferase activity expressed by *C. parvum* sporozoites collected from the duodenal (Duo), jejunal (Jej) or ileal (Ile) TIM-1 compartments during child (purple) or adult (dark blue) digestion. Boxplots depict the relative luminescence of sporozoites compared with that of the inoculum for each digestion assay. For each age condition, significant differences (*P* < 0.05) between time points are indicated by non-corresponding letters (purple lowercase for child digestion and blue capitals for adult digestion). For each time point, significant differences between age conditions are indicated in black as follows: **** *P* < 0.0001. **(B)** Invasion assay. *C. parvum* sporozoites were allowed to infect HCT-8 monolayers for 2.5 h before evaluation of luminescence intensity. Boxplots depict the luminescence measured in infected cells from which the luciferase background measured in non-infected cells has been removed. For each age condition, significant differences (*P* < 0.01 or *P* < 0.001 for child or adult, respectively) between time points are indicated by non-corresponding letters (purple lowercase for child digestion and blue capitals for adult digestion). For each time point, significant differences between age conditions are indicated in black as follows: ** *P* < 0. 01.

The invasion ability of Nluc-expressing sporozoites collected and purified from the TIM-1 small intestine was assessed by *in vitro* infection of HCT-8 cell monolayers and subsequent determination of luciferase activity (Figure 3.B). Time of digestion had an overall significant effect on sporozoite ability to invade host cells in both child (*P* < 0.0001) and adult (*P* < 0.0001) conditions. The invasion of HCT-8 monolayers was observed to be low and stable for sporozoites collected from the duodenum at 60 min or from the jejunum at 120 min of both child (9.3 × 10^2^ ± 3.4 × 10^2^ RLUs and 1.4 × 10^3^ ± 2.5 × 10^2^ RLUs, respectively) and adult (3. × 10^2^ ± 1.2 × 10^2^ RLUs and 4.6 × 10^2^ ± 1.6 × 10^2^ RLUs, respectively) digestive conditions. The invasion ability was observed to be significantly increased for sporozoites collected at 180 min from the child ileum (8.0 × 10^3^ ± 3.0 × 10^3^ RLUs; *P* < 0.01 between 180 min and other time points) or the adult ileum (4.8 × 10^3^ ± 8.0 × 10^2^ RLUs; *P* < 0.0001 between 180 min and other time points). By the end of digestion, parasite invasion dropped to a level comparable to the one observed at 60 min or 120 min, with a relative luminescence of 1.0 × 10^3^ ± 2.8 × 10^2^ or 5.1 × 10^2^ ± 1.3 × 10^2^ RLUs detected for monolayers infected with sporozoites collected at 300 min in the child or adult ileum, respectively. The invasion ability of sporozoites was significantly higher in children compared with adults only at 120 min in the jejunum (*P* < 0.01).

### Parasite gene expression following child or adult digestion in the TIM-1 model

The parasite transcriptome modifications induced by adult or child digestions in the TIM-1 were determined by whole transcriptome sequencing (RNA-Seq) for gastric (0-20 min) and ileal (120-180 min) effluents, and also for the initial inoculum as control. A principal component analysis (PCA) was performed to evaluate the samples’ distribution according to their expression profiles. The first (31%) and second (16%) components represented most of the differential expression pattern with a cumulative proportion of 47% (Figure 4.A). This PCA analysis revealed a segregation between samples collected at different time points and from different TIM-1 compartments. The PCA plot showed that all six samples from the ileal effluents were grouped together, and that the three inoculum samples and the six samples from the gastric effluents formed two other distinct groups (*i.e.,* clusters), despite samples ‘Inoculum_3’ and ‘Stomach_Adult2’ being more distant from their respective replicates. For each cluster corresponding to samples collected from gastric or ileal effluents, no further segregation by age could be observed, suggesting that most of the variance is explained by time and/or digestive compartment, rather than simulated age (*i.e.,* child *vs.* adult). Indeed, pairwise comparisons resulted in no or only one gene (Gene ID CPATCC_0017110: unspecified product) whose expression was significantly modified (*i.e.,* showing an adjusted *P*-value [FDR] < 0.05 and a minimum 2-fold regulation [log2FC > 1]) between the two age conditions when effluent was collected in the gastric or in the ileal compartment, respectively (Figure 4.B, Additional files 2 and 3). In contrast, expression profiles were more affected by time and/or digestive compartment, with for example 22 genes whose transcription levels were modified between gastric samples and inoculum for both age conditions, and 83 or 99 genes between gastric and ileal samples for child or adult conditions, respectively. Furthermore, the strongest modulation of the parasite transcriptome was observed between ileal samples and inoculum, with 190 genes whose transcription levels were significantly affected in the adult digestive condition and up to 208 genes in the child.

**FIGURE 4.**
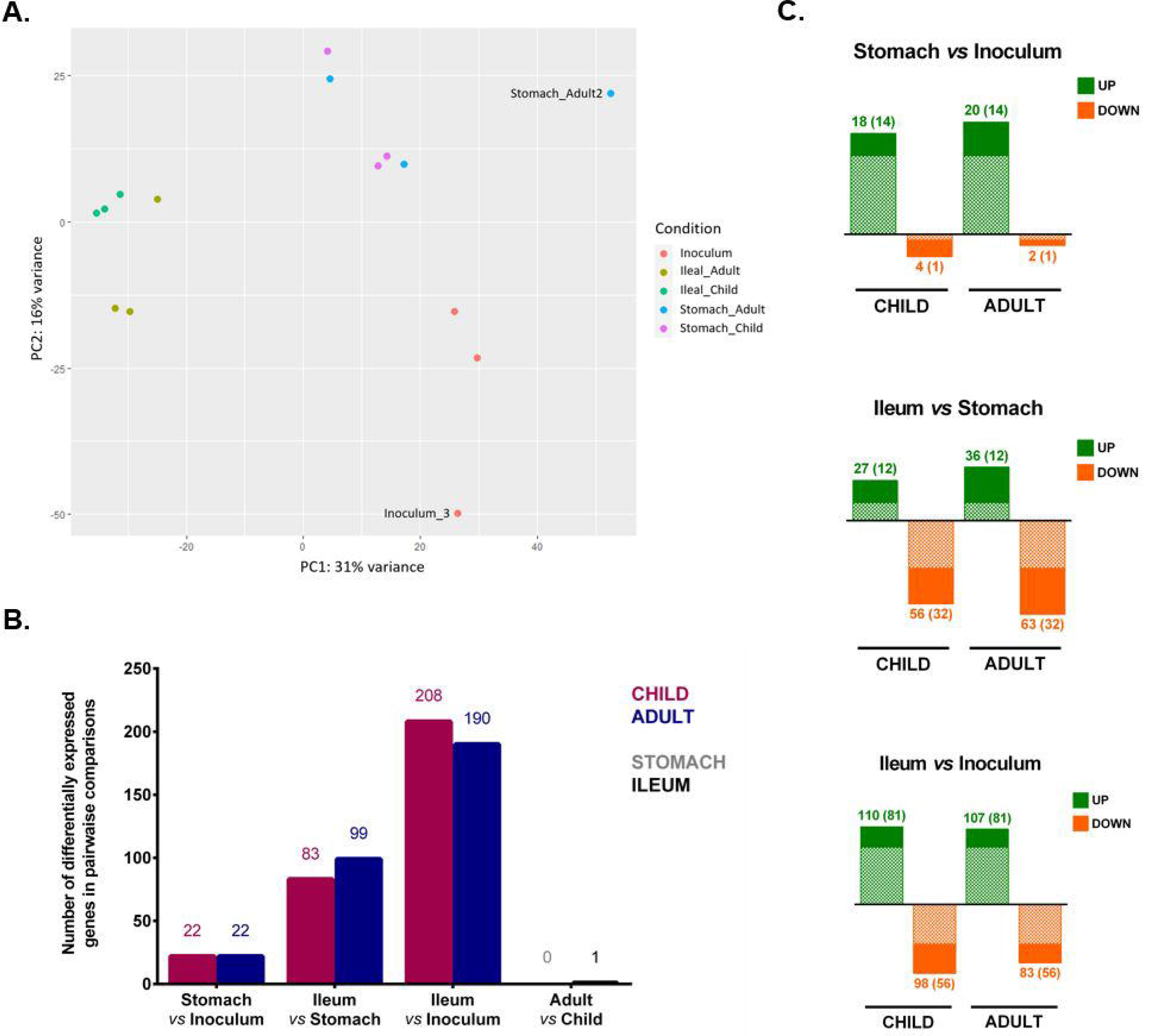
RNA-Seq analysis of *C. parvum* genes in TIM-1 samples. **(A)** Principal component analysis of RNA-Seq samples investigating gene expression changes in *C. parvum* parasites collected in the inoculum (t = 0 min), or in gastric (0-20 min) or ileal effluents (120-180 min) during the simulated child or adult digestions in the TIM-1. PCA was performed using normalised RNA-Seq data of a set of 2430 filtered genes. **(B)** Number of differentially expressed genes (DEGs) between pairwise comparisons. All DEGs show an adjusted *P*-value < 0.05 and a minimal regulation of 2-fold (log2FC > 1). **(C)** Number of upregulated (green) or downregulated (orange) genes between samples collected from different TIM-1 compartments in both child and adult conditions. The number of DEGs shared by age conditions is shown in brackets and with the dotted filling pattern.

Most of the differentially expressed genes (DEGs) detected in samples collected from the stomach between 0 and 20 min and compared to the inoculum were found upregulated: 18/22 or 20/22 for child or adult, respectively, with 14 genes shared by both age conditions (Figure 4.C, Additional files 4 and 5). An analysis of enriched gene ontology (GO) performed on these overexpressed genes in the stomach of the child condition found two significantly enriched GO terms related to cellular components (Additional files 4 and 6.A): ribonucleoprotein complex (FDR = 0.026) and ribosome (FDR = 0.026), both associated with the translation process. Approximately two thirds (*i.e.,* 67.4% or 63.6% for child and adult conditions, respectively) of the DEGs identified in ileal samples compared to gastric samples were found downregulated (Figure 4.C, Additional files 7 and 8). Thirty-two significantly downregulated genes are shared by child and adult conditions, among which a gene encoding a myosin motor domain containing protein (Gene ID CPATCC_0009620) showed a 6-fold decrease. One gene encoding an actin protein (Gene ID CPATCC_0025920) showed a 3.8-fold decrease, but only in the child condition. Among downregulated genes, GO enrichment analysis revealed a single significantly enriched GO term related to a molecular function in the adult condition: protein tyrosine/serine/threonine phosphatase activity (FDR = 0.024). Six genes that were found significantly upregulated in the child ileum compared to the child stomach encode putative secreted proteins, of which two are part of the SKSR family (Gene IDs CPATCC_0030860 and CPATCC_0035410) and one is an alpha/beta hydrolase (Gene ID CPATCC_0025460), the latter being also significantly upregulated in the adult condition. The GO enrichment analysis found six significantly enriched GO terms in the child (FDR < 0.05), all related to diverse cellular components (*i.e.,* nuclear or organelle compartments) (Additional files 6.B and 7). The expression of a gene encoding an actin family protein showed a 40-fold significant increase in the adult condition (Gene ID CPATCC_0036850).

As mentioned above, the strongest modification of the parasite transcriptome was observed for ileal samples between 120 and 180 min of digestion when compared to the initial inoculum (Figure 4.B, Additional files 9 and 10). For instance, 98 and 83 DEGs were significantly downregulated in child and adult ileum, respectively, with 56 genes shared by both age conditions (Figure 4.C). Among shared downregulated genes, the expression of the gene CPATCC_0009860 encoding the cryptosporidial mucin, also designated as glycoprotein-900 (GP900), was significantly decreased by a 4.8-fold. The myosin motor domain containing protein-encoding gene CPATCC_0009620 was also significantly downregulated in the ileum compared to the inoculum (9.7- or 6.8-fold decrease in the child or the adult condition, respectively). Subsequent metabolic pathway enrichment analysis interrogating the KEGG pathway database with significantly downregulated genes highlighted glycolysis/gluconeogenesis as the single significantly enriched pathway in the child ileum (FDR = 0.029) (Figure 5.A). The expression of 110 and 107 genes was significantly increased in child and adult ileum, respectively, among which the majority (*i.e.,* 81 genes) was shared by both age conditions (Figure 4.C). For example, four genes encoding putative secreted protein were significantly upregulated in both child and adult ileum, when compared to the inoculum. Among those genes, one relates to the putative secreted alpha/beta hydrolase previously identified (Gene ID CPATCC_0025460) and two belong to the SKSR gene family (CPATCC_0000030 and CPATCC_0030860). GO enrichment analysis performed on significantly upregulated genes highlighted 22 significantly enriched GO terms (FDR < 0.05) shared by child and adult (Figure 5, Additional files 9 and 10). Among these, the top three shared enriched GO terms were associated with cellular components and identified as intracellular structure, intracellular organelle and organelle. Gene expression was shared by both age conditions and associated to 26 or 32 genes out of the 110 (23.6%) or 107 (29.9%) significantly upregulated genes detected in the child or the adult condition, respectively. Using the KEGG pathway database, folate biosynthesis was identified as the top one enriched pathway in child (FDR = 0.0008) or adult (FDR = 0.010) ileum compared to the inoculum, and was mostly associated with upregulated helicase-encoding genes. Interestingly, the expression of one gene associated to myosin complex and cytoskeletal motor activity (Gene ID CPATCC_0021030), was highly significantly upregulated by a 1978-fold in the child ileum, compared to the inoculum sample.

**FIGURE 5.**
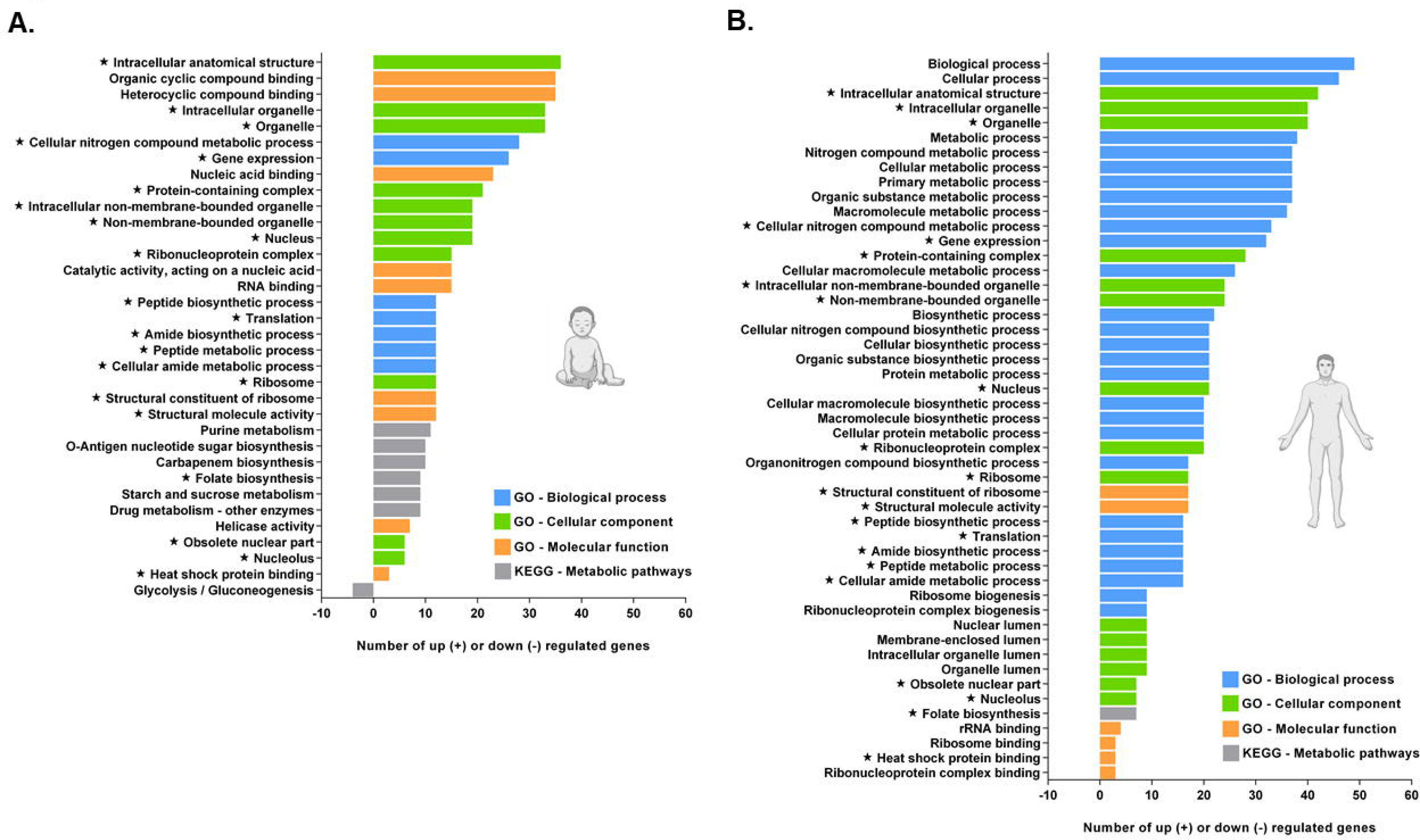
Enrichment analysis on *C. parvum* genes differentially expressed in the ileal compartment. The Gene Ontology and KEGG enrichment analysis was performed on genes that were significantly up- or downregulated in the ileal effluents compared with the inoculum. The number of genes associated with each significantly enriched (*i.e.,* showing a Benjamini-Hochberg-adjusted *P*-value or FDR < 0.05) GO term or KEGG pathway is displayed for the child **(A)** or the adult **(B)** conditions. The three GO term categories named ‘Biological process’, ‘Cellular component’ and ‘Molecular function’ are represented by blue, green or orange bars, respectively. The KEGG Metabolic pathways are represented by grey bars. The GO terms or KEGG pathway shared by both age conditions are indicated by a star.

## DISCUSSION

The goal of our study was to perform a comparative analysis of *C. parvum* infection under the digestive conditions encountered in young children (from 6 months to 2 years) *versus* those in adults. Due to the inherent limitations of *in vivo* experimentation within the host, the investigation of *C. parvum* excystation has only been performed to date with highly simplified *in vitro* approaches integrating only a few digestive parameters simultaneously (*e.g.*, temperature, pH) (29). Furthermore, these approaches have never been conducted under finely defined child *vs.* adult digestive conditions, nor combining digestive physicochemical properties with transit dynamism. Based on previous studies on other pathogens (10,12–14,16), we used the dynamic multicompartmental TIM-1 system as a suitable model to monitor *C. parvum* excystation along the simulated human upper gastrointestinal tract of both infant and adult conditions. Our study demonstrates for the first time that most oocysts are found intact in the simulated human stomach compartment 45 min after inoculation in the TIM-1 and that the excystation step occurs rapidly and is almost completed within the following 15 min and upon passage in the duodenal compartment. These data suggest that the majority of *C. parvum* oocysts do not undergo excystation until reaching the duodenum, thereby avoiding premature release and subsequent inactivation of the more susceptible sporozoite stages in the gastric acid environment. This is in accordance with the existence of two *Cryptosporidium* lineages that show adaptation to different excystation conditions, one displaying gastric tropism including parasite species that multiply in this gastric acidic environment (*e.g.*, *C. andersoni* and *C. muris*) and one exhibiting intestinal tropism (*e.g.*, *C. parvum* and *C. hominis*) (30). In order to maximise the delivery of sporozoites according to species tropism, excystation is thus activated by different host-derived triggers. In the TIM-1 system, mirroring *in vivo* situations, the excystation of *C. parvum* oocysts might have been triggered first by elevation of the temperature to 37°C upon parasite inoculation into the system, followed by a drastic change in pH upon passage from the acid gastric compartment to the alkaline duodenal one. Previous *in vitro* studies investigating host-derived factors triggering *C. parvum* oocysts excystation have reported that temperature increase and pH change play the most important roles in the transduction of external signals across the oocyst wall to sporozoites (29,31–33). The digestive secretions (*i.e.,* bile salts, trypsin and pancreatic juice) that the duodenal compartment receives in the TIM-1 system might also play a role in *C. parvum* excystation process, although our knowledge of their precise role as excystation triggers or putative synergistic effect is still limited. Interestingly, *in vitro* incubation of oocysts in bile, in particular in sodium taurocholate following exposure to an acid pre-treatment, has been shown to enhance parasite excystation, mimicking transition from the acidic gastric environment to the alkaline small intestine (29,31,34).

Our flow cytometry analysis suggests that the excystation rate was slightly more efficient along the simulated small intestine compartments upon adult digestive condition compared to the child one. This could be attributable to specific differences in the physicochemical parameters described for child *versus* adult physiology in a healthy state, which were subsequently implemented into the TIM-1 programs (12). For instance, the discrepancy in excystation success may be explained by the slower and less pronounced gastric pH acidification in children (35,36) and/or by the lower concentrations of various secretion components delivered in the child’s duodenum, especially of bile salts (37,38) known to enhance excystation (29,31,34).

The constitutively expressed luciferase marker was used as an indicator to assess sporozoite physiological state, since the intensity of its robust and sensitive signal correlates directly with the number and viability of parasites (Additional file 1). Taken together, our flow cytometry and luciferase analyses suggest that the highest amount of released sporozoites is present in the simulated ileum, and their physiological state peaks in this compartment at 180 min of digestion, regardless of age conditions. This suggests that *C. parvum* has evolved an excystation process that ensures the accumulation of the maximum number of freshly released sporozoites in the gastrointestinal segment where this species shows the highest tissue tropism *in vivo*. While *C. parvum* has been reported to colonise both proximal intestinal segments and the colon in different hosts (39,40), infections associated with this species are predominantly concentrated in the distal small intestine (41,42). For instance, recent *in vivo* bioluminescent imaging of a Nluc-expressing *C. parvum* strain throughout the intestinal tract of IFN-γ-KO mice clearly showed the parasites to be mainly localised in the ileum section as well as in the caecum (43).

Our results also suggest that simulated small intestine of the child condition is associated with fewer sporozoites on the one hand, but a significantly higher sporozoite luciferase activity on the other hand compared to the adult condition. Consequently, sporozoites residing in the intestinal tract of infants could be characterised by a higher viability or be in a less damaged state. In the context of a contamination by the major food- or waterborne pathogen enterohemorrhagic *Escherichia coli* (EHEC O157:H7 serotype), Roussel *et al.* used a similar TIM-1 setup to demonstrate that a higher amount of viable bacterial cells may reach the distal parts of the child’s small intestine, compared to those of adults (12). These results might be attributed to less stringent conditions encountered by bacteria or *C. parvum* sporozoites in the child’s intestinal tract, in regard to the two-fold lower concentration in bile salts and pancreatic secretions described for children compared to adults (37,38,44,45), also reproduced in the TIM-1 model. To our knowledge, the direct impact of various concentrations of duodenal secretions on *C. parvum* sporozoite survival has not yet been investigated. Therefore, we can not exclude that the significantly decreased physiological state observed for the sporozoites released in the adult small intestine may be the result of the detergent properties associated with higher concentrations of duodenal secretions encountered in this environment.

We then aimed to investigate whether the higher physiological state of *C. parvum* sporozoites observed in the child intestinal tract could also correlate with an increased infectivity. To this end, the TIM-1 system was combined for the first time with a cell culture assay, wherein we tested the capacity of sporozoites collected from the digestive compartments to invade human intestinal epithelial cells. The invasion assay suggests that sporozoites collected from the ileal compartment at 180 min of digestion have a significantly higher ability to invade HCT-8 cells, compared to other time points. This finding may reflect the slightly higher amount of sporozoites present in the ileum compartment, as compared to the duodenum compartment for instance, but it may also be linked to the significantly higher sporozoites’ physiological state and further emphases the tropism of *C. parvum* for the ileum *in vivo*. Interestingly, when collected from the same TIM-1 compartment, but at the end of the digestion process (300 min of digestion), sporozoites show a decreased luciferase activity and demonstrate a markedly reduced ability to invade host cells. In the literature, *C. parvum* sporozoites have been reported as somewhat fragile (46–48), surviving only for a few hours after release from the much more resilient oocyst stage. Our results are in agreement with this description and suggest that the infectivity potential of these vulnerable parasite stages may decrease drastically in the small intestine lumen if invasion of epithelial cells does not occur quickly after excystation. Our invasion assay shows that, although a lower amount of sporozoites was inoculated to host cells under the child condition (*i.e.,* according to our flow cytometry data), these sporozoites exhibit equivalent or even significantly higher invasion ability compared to the adult condition, most likely due to their “better” physiological state. Thus, our results suggest that age-related variability in digestive physicochemical parameters may modulate different features of sporozoite physiology, and therefore participate in higher susceptibility of young children to *C. parvum* infection compared to adults.

In this study, we also aimed to characterise the modulation of the *C. parvum* gene expression in response to the various digestive physicochemical parameters encountered by the parasite in two different compartments of the child or adult gastrointestinal tract. Intriguingly, our results revealed that the parasite transcriptome is almost exclusively affected by the time and/or compartment of digestion in the TIM-1, rather than by the simulated age (*i.e.,* child *vs.* adult). This observation suggests that parasite gene expression is predominantly governed by the succession of shifts in digestive environmental conditions and the duration of digestion, rather than by more subtle physicochemical specificities associated with age condition. In contrast, previous RT-qPCR analyses performed on TIM-1 gastric and ileal effluents have shown that the expression of major EHEC O157:H7 bacterial virulence genes was significantly higher under child digestive conditions, compared to the adult ones (12). Regarding *C. parvum*, although differences in digestive physicochemical properties between infants and adults may influence the amount, physiological state and invasiveness of sporozoites released in the small intestine, they may not be associated with significant modulation of parasite genes associated with virulence.

Transcriptomic analysis performed on TIM-1 gastric effluents collected within the initial twenty minutes of digestion detected a low number of DEGs between stomach and inoculum samples, predominantly exhibiting upregulation. In this digestive compartment, where most parasites were found as intact oocysts, enrichment of upregulated genes associated with gene expression and translation was detected. Similarly, previous transcriptomic analysis has revealed that the *C. parvum* oocyst stage is highly active in protein synthesis, as evidenced by high transcripts levels of parasite genes involved in ribosome biogenesis, transcription and translation (49).

We also analysed the *C. parvum* transcriptome collected from TIM-1 ileal effluents between 120 and 180 min of digestion, when the vast majority of parasites are found as released sporozoites compared to the stomach compartment. Our RNA-Seq analysis revealed that the sporozoite-enriched transcriptome exhibited much greater modulation in the ileum compared to the stomach, with approximately 25% of upregulated genes associated with gene expression, nucleus or intracellular structures and organelles. KEGG analysis highlighted the glycolysis/gluconeogenesis as a significantly downregulated pathway in parasites collected from ileal effluents. Although *C. parvum* may possess a remnant mitochondrion, it lacks the Krebs cycle and the cytochrome-based respiration, therefore relying mainly, if not only, on glycolysis for ATP production (50). Glycolysis-related genes are known to be highly expressed in intracellular developmental stages (51), supporting *C. parvum* replication inside host cells. While some of their related products have also been detected in the sporozoite proteome (52), they were found under-represented in this repertoire in another proteomic study (53).

Previous transcriptomic analysis has shown that, in contrast to intracellular developmental stages, *C. parvum* extracellular stages (*i.e.,* oocysts and sporozoites) express a wider range of genes encoding specialised functions (54), with few identified orthologs outside or related protozoan organisms. Like other apicomplexan parasites, the polarised *C. parvum* sporozoites harbour unique apical secretory organelles involved in attachment, invasion, penetration and maintenance of the parasite within the host cell, (*i.e.*, micronemes, dense granules, small granules and a single rhoptry), as well as the canonical glideosome dependent on actin and myosin providing parasite gliding motility (55–57). Accordingly, the expression of several parasite genes associated with the latter apicomplexan feature, particularly the myosin complex and cytoskeletal motor activity, was significantly modulated in the sporozoite-enriched fractions collected from the TIM-1 ileal effluents. While two of these genes (Gene IDs CPATCC_0009620 and CPATCC_0025920) showed a moderate downregulation in these samples, two other encoding a protein from the actin family (Gene ID CPATCC_0036850) and a myosin motor domain containing protein (Gene ID CPATCC_0021030) were characterised by a tremendous 40- or ∼ 2000-fold upregulation, respectively. Alongside this marked regulation of genes potentially related to parasite gliding motility, the expression of genes encoding putative secreted proteins was significantly modulated in ileal effluents. For instance, the expression levels of the mucin-like glycoprotein GP900 (Gene ID CPATCC_0009860) were significantly downregulated in our ileal sporozoite-enriched samples. Stored in the sporozoite micronemes and previously hypothesised to be involved in attachment and/or invasion (58), the immunodominant protein GP900 has recently been shown to enter the secretory pathway after excystation, where its short cytoplasmic domain is cleaved before discharge of the cleaved form to the extracellular space, suggesting a lubrication role during sporozoite invasion (59). Finally, six genes encoding putative secreted proteins were significantly upregulated in the TIM-1 ileal effluents, consistent with the past detection of several proteins associated with extracellular protein secretion in the protein repertoire of *C. parvum* sporozoites (52,53). Apart from a predicted hydrolase, the function of these putative secreted proteins is not yet known. Interestingly, three of these genes are predicted to encode proteins belonging to the SKSR family (Gene IDs CPATCC_0030860, CPATCC_0000030 and CPATCC_0035410). This *Cryptosporidium*-specific multigene family comprises most highly polymorphic and subtelomeric genes encoding secreted proteins harbouring a signal peptide and SK and SR repeats (60). Although present in all major human-infecting *Cryptosporidium* species, differences in the presence or absence, as well as in the copy numbers of this subtelomeric gene family, were recently identified between sequenced *Cryptosporidium* genomes by comparative genomic analyses, similarly to two other families encoding the MEDLE secretory proteins, named after their conserved sequence motif at the C-terminus, and insulinase-like proteases (61–66). Recent findings identified SKSR1 as a member of the secretory proteins (67) in the newly identified small granules organelles (57). Additionally, SKSR1 was shown to be secreted into the parasite-host interface (*i.e.*, parasitophorous vacuole membrane and feeder organelle) and to be important for *C. parvum* pathogenicity, suggesting that it may act as a virulence factor through regulating host responses (67). More comparative studies are needed to fully elucidate the function of all SKSR members, since gene gains and losses of this subtelomeric gene family are suggested to contribute to differences in pathogenicity and host specificity in *Cryptosporidium* populations.

## CONCLUSIONS

Being one of the most common causes of infectious moderate-to-severe diarrhoea in young children, cryptosporidiosis is an important contributor to early childhood mortality in low-resource settings. Increased susceptibility of infants and toddlers to *Cryptosporidium* can be attributed to immature immune status in this age group. Using the sophisticated TIM-1 *in vitro* gastrointestinal model, we showed that the digestive physicochemical parameters encountered in the child digestive tract could be associated with fewer, however potentially more active and invasive sporozoites released. Our results suggest that age-mediated variation in the human gastrointestinal physiology could also partially explain why young individuals are more at risk. The link between specificities of the child digestive tract and disease is complex. Understanding the interactions between *C. parvum* infection and various digestive components of young individuals, including physicochemical parameters, mucus barrier, and microbiota, can provide us with a more accurate picture of children susceptibility, and possible valuable information towards new treatment strategies.

## LIST OF ABBREVIATIONS

ATP: Adenosine triphosphate
*C. andersoni*: *Cryptosporidium andersoni*
*C. hominis*: *Cryptosporidium hominis*
*C. muris*: *Cryptosporidium muris*
*C. parvum*: *Cryptosporidium parvum*
DEG: Differentially expressed gene
DNA: Deoxyribonucleic acid
DNase: Deoxyribonuclease
*E. coli*: *Escherichia coli*
EHEC O157:H7: Enterohemorrhagic *Escherichia coli* serotype O157:H7
FBS: Fetal bovine serum
FDR: False discovery rate
GO: Gene ontology
GP900: Glycoprotein-900
IFN-γ: Interferon gamma
KEGG: Kyoto encyclopedia of genes and genomes
KO: Knock-out
Nluc: Nanoluciferase
PBS: Phosphate-buffered saline
PCA: Principal component analysis
RLU: Relative light unit
RNA: Ribonucleic acid
RNA-Seq: RNA sequencing
RPMI: Roswell Park Memorial Institute
SEM: Standard error of the mean
TIM-1: TNO (Toegepast Natuurwetenschappelijk Onderzoek) gastrointestinal model-1

## DECLARATIONS

### Ethics approval and consent to participate

Not applicable

### Consent for publication

Not applicable

### Availability of data and materials

The datasets generated and analysed during the current study are available in the GEO repository database, under accession number GSE271211 (https://www.ncbi.nlm.nih.gov/geo/query/acc.cgi?&acc=GSE271211).

### Competing interests

The authors declare that they have no competing interests.

## Funding

This research was supported by the INRAE Animal Health division and by the Laboratoire d’Excellence (LabEx) ParaFrap (ANR-11-LABX-0024).

## Author’s contributions

JT, LE-M, SB-D and SL-L designed the project and the experiments. JT, LE-M, SC, AS, SD, CM and SL-L performed the experiments. JT and AS produced and purified the oocysts of the *C. parvum* INRAE Nluc strain. LE-M, SC, SD and CM performed the *in vitro* digestions. SL-L performed the flow cytometry analysis with the help of CB. JT performed the sporozoite luciferase activity and the cell culture experiments. AS performed the parasite RNA extractions. JT and GS analysed the RNA-Seq data. JT, LE-M, GS, FL, SB-D and SL-L interpreted the experimental work. JT, LE-M, FL, SB-D and SL-L secured funding. JT, LE-M, SB-D and SL-L wrote the paper with editorial support and comments from all other authors. All authors read and approved the final version of the manuscript.

## Acknowledgments

We would like to thank the Helixio company for the mRNA sequencing and bioinformatics analysis. We are also very grateful to Elise Courtot (UMR ISP, INRAE) for her help on statistical analysis.

---

**ADDITIONAL FILE 1**

**File format.** Figure (.pdf)

**Title of data.** Luciferase activity expressed by live or heat-killed *C. parvum* INRAE Nluc parasites.

**Description of data.** Luciferase activity expressed by live or heat-killed *C. parvum* INRAE Nluc parasites. Luciferase activity was measured in live (*i.e.*, maintained at room temperature for 30 min, blue dots) or heat-killed (*i.e.*, maintained at 60°C for 30 min, red dots) *C. parvum* INRAE Nluc parasites, following excystation of 1 × 10^7^ oocysts or 1 × 10^6^ oocysts. Individual values from five independent experiments are represented. Significant differences between groups determined by Mann-Whitney non-parametric test are indicated as follows: ** *P* < 0.01. Cp, *Cryptosporidium parvum*. RT, room temperature.

**ADDITIONAL FILE 2**

**File format.** Excel spreadsheet (.xls)

**Title of data.** RNA-Seq Stomach Adult *vs* Stomach Child.

**Description of data.** Differentially expressed genes determined by RNA-sequencing between samples collected in gastric effluents of adult and gastric effluents of child, and results from enrichment analyses performed on significantly upregulated or downregulated genes.

**ADDITIONAL FILE 3**

**File format.** Excel spreadsheet (.xls)

**Title of data.** RNA-Seq Ileal Adult *vs* Ileal Child.

**Description of data.** Differentially expressed genes determined by RNA-sequencing between samples collected in ileal effluents of adult and ileal effluents of child, and results from enrichment analyses performed on significantly upregulated or downregulated genes.

**ADDITIONAL FILE 4**

**File format.** Excel spreadsheet (.xls)

**Title of data.** RNA-Seq Stomach Child *vs* Inoculum.

**Description of data.** Differentially expressed genes determined by RNA-sequencing between samples collected in gastric effluents of child and inoculum, and results from enrichment analyses performed on significantly upregulated or downregulated genes.

**ADDITIONAL FILE 5**

**File format.** Excel spreadsheet (.xls)

**Title of data.** RNA-Seq Stomach Adult *vs* Inoculum.

**Description of data.** Differentially expressed genes determined by RNA-sequencing between samples collected in gastric effluents of adult and inoculum, and results from enrichment analyses performed on significantly upregulated or downregulated genes.

**ADDITIONAL FILE 6**

**File format.** Figure (.pdf)

**Title of data.** Enrichment analysis on *C. parvum* genes differentially expressed in the stomach compared to the inoculum **(A)** or in the ileum compared to the stomach **(B)**.

**Description of data.** The Gene Ontology and KEGG enrichment analysis was performed on genes that were significantly up- or downregulated. The number of genes associated with each significantly enriched (*i.e.,* showing a Benjamini-Hochberg-adjusted *P*-value or FDR < 0.05) GO term or KEGG pathway is displayed for both child (left) and adult (right) conditions. The three GO term categories named ‘Biological process’, ‘Cellular component’ and ‘Molecular function’ are represented by blue, green or orange bars, respectively. The KEGG Metabolic pathways are represented by grey bars.

**ADDITIONAL FILE 7**

**File format.** Excel spreadsheet (.xls)

**Title of data.** RNA-Seq Ileal Child *vs* Stomach Child.

**Description of data.** Differentially expressed genes determined by RNA-sequencing between samples collected in ileal effluents of child and gastric effluents of child, and results from enrichment analyses performed on significantly upregulated or downregulated genes.

**ADDITIONAL FILE 8**

**File format.** Excel spreadsheet (.xls)

**Title of data.** RNA-Seq Ileal Adult *vs* Stomach Adult.

**Description of data.** Differentially expressed genes determined by RNA-sequencing between samples collected in ileal effluents of adult and gastric effluents of adult, and results from enrichment analyses performed on significantly upregulated or downregulated genes.

**ADDITIONAL FILE 9**

**File format.** Excel spreadsheet (.xls)

**Title of data.** RNA-Seq Ileal Child *vs* Inoculum.

**Description of data.** Differentially expressed genes determined by RNA-sequencing between samples collected in ileal effluents of child and inoculum, and results from enrichment analyses performed on significantly upregulated or downregulated genes.

**ADDITIONAL FILE 10**

**File format.** Excel spreadsheet (.xls)

**Title of data.** RNA-Seq Ileal Adult *vs* Inoculum.

**Description of data.** Differentially expressed genes determined by RNA-sequencing between samples collected in ileal effluents of adult and inoculum, and results from enrichment analyses performed on significantly upregulated or downregulated genes.

## Notes

https://www.ncbi.nlm.nih.gov/geo/query/acc.cgi?&acc=GSE271211

### Competing Interest Statement

The authors have declared no competing interest.

