## Supplementary figures and images for "Exploring the impact of digestive physicochemical parameters of adults and infants on the pathophysiology of *Cryptosporidium parvum* using the dynamic TIM-1 gastrointestinal model"

### Additional File 1

Additional file 1

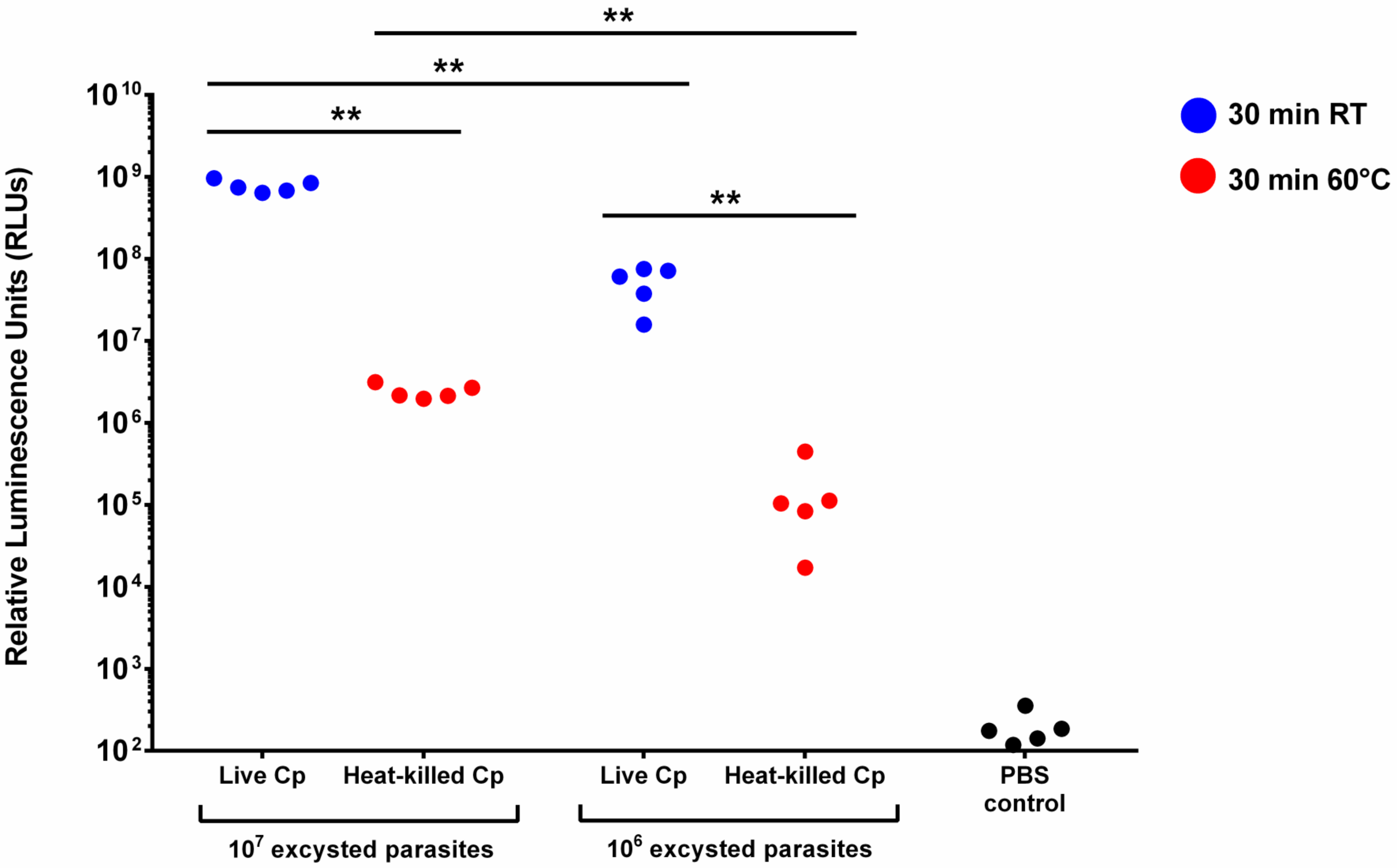

### Additional File 6

Additional file 6

**A.**

STOMACH *vs.* INOCULUM

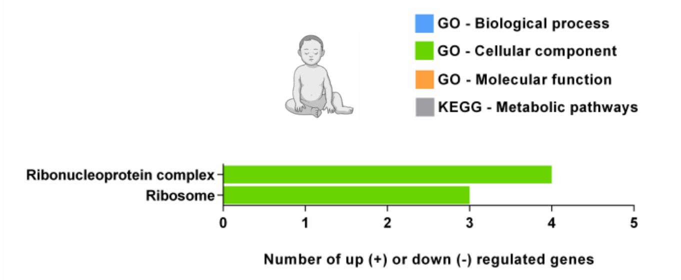


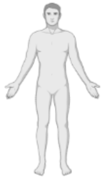

NONE

**B.**

ILEUM *vs.* STOMACH

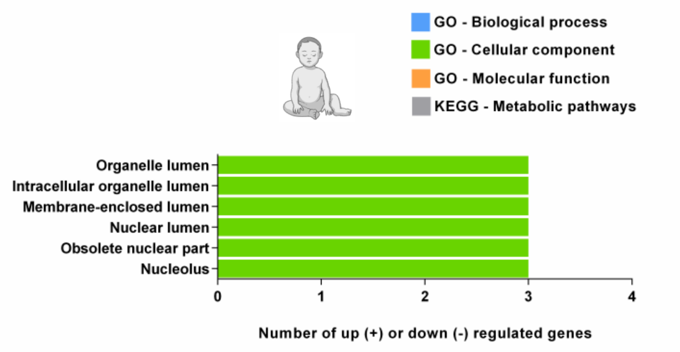


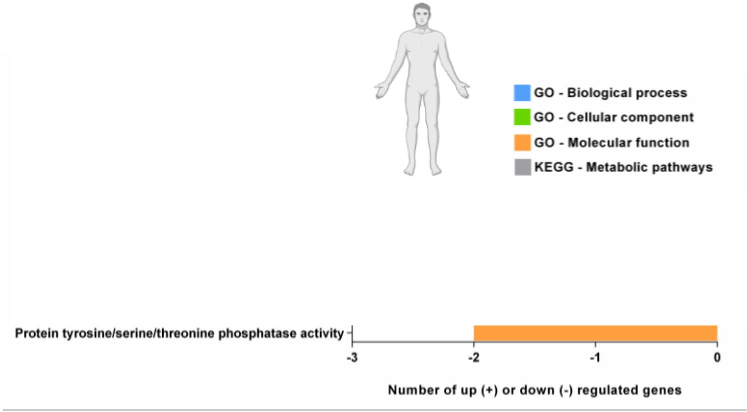
